# Design and optimization of a cell-free atrazine biosensor

**DOI:** 10.1101/779827

**Authors:** Adam D. Silverman, Umut Akova, Khalid K. Alam, Michael C. Jewett, Julius B. Lucks

**Affiliations:** Department of Chemical and Biological Engineering, Northwestern University, Evanston, IL 60208, USA; Chemistry of Life Processes Institute, Northwestern University, Evanston, IL 60208, USA; Center for Synthetic Biology, Northwestern University, Evanston, IL 60208, USA; Interdisciplinary Biological Sciences Program, Northwestern University, Evanston, IL 60208; Member, Robert H. Lurie Comprehensive Cancer Center, Northwestern University, Chicago, IL 60611, USA; Member, Simpson Querrey Institute, Northwestern University, Chicago, IL 60611, USA; Center for Water Research, Northwestern University, Evanston, IL 60208, USA

**Keywords:** cell-free, TX-TL, metabolism, transcription factor, biosensor, atrazine, cyanuric acid, synthetic biology

## Abstract

Recent advances in cell-free synthetic biology have spurred the development of *in vitro* molecular diagnostics that serve as effective alternatives to whole-cell biosensors. However, cell-free sensors for detecting manmade organic water contaminants such as pesticides are sparse, partially because few characterized natural biological sensors can directly detect such pollutants. Here, we present a platform for the cell-free detection of one critical water contaminant, atrazine, by combining a previously characterized cyanuric acid biosensor with a reconstituted atrazine-to-cyanuric acid metabolic pathway composed of several protein-enriched bacterial extracts mixed in a one pot reaction. Our cell-free sensor detects atrazine within an hour of incubation at an activation ratio superior to previously reported whole-cell atrazine sensors. We also show that the response characteristics of the atrazine sensor can be tuned by manipulating the component ratios of the cell-free reaction mixture. Our approach of utilizing multiple metabolic steps, encoded in protein-enriched cell-free extracts, to convert a target of interest into a molecule that can be sensed by a transcription factor is modularly designed, which should enable this work to serve as an effective proof-of-concept for rapid field-deployable detection of complex organic water contaminants.

## INTRODUCTION

Cell-free gene expression (CFE) has recently emerged as a powerful strategy for rapid, field-deployable diagnostics for nucleic acids^1–5^ and chemical contaminants.^6–9^ One reason for this success is that CFE reactions minimize many of the constraints of whole-cell sensors, including mass transfer barriers, cytotoxicity, genetic instability, plasmid loss, and the need for biocontainment.^8,10^ In addition, CFE reactions can be stabilized through freeze-drying and then are activated upon rehydration, enabling the biosensors to be used outside the laboratory at the point of sampling in the field.^1^ However, previously reported cell-free biosensors have so far been limited to detecting either nucleic acids^2,3^ or chemical contaminants that can be directly sensed with well-characterized allosteric protein transcription factors or riboswitches.^8,9,11–13^

Here, we expand the ability of cell-free biosensors to detect complex organic molecules by developing a combined metabolism and biosensing strategy. Our strategy is motivated by the observation that the space of known natural transcription factors may be insufficient to directly detect organic molecules of analytical interest, especially those that are man-made and relatively new to natural environments. On the other hand, a wealth of metabolic biochemistry often exists that could convert a target molecule of interest into a compound that can be directly sensed by a transcription factor. Thus, a range of new CFE-based diagnostics could be developed by combining *in vitro* metabolic conversion with natural transcriptional biosensors. Recently, such a strategy was validated in CFE reactions using a simple one-enzyme pathway where the enzyme, transcription factor, and reporter are encoded on separate plasmids. In that work, cocaine and hippuric acid were catabolized *in vitro* to make benzoic acid, which is sensed by the allosteric transcription factor BenR.^7,14,15^ However, the metabolic pathways tested were short – containing only a single enzyme – and converged to a simple and abundant analyte detected by a native *E. coli* transcription factor. A more general approach would be necessary for detecting xenobiotic, or new-to-nature, analytes.

Specifically, in this study, we develop a strategy for multi-enzymatic metabolic biosensing of atrazine–one of the most commonly detected herbicides in American surface water, and a suspected endocrine-disrupting compound.^16^ Atrazine is frequently measured in finished water sources at concentrations above the recommended 3 parts per billion (ppb) Maximum Containment Level Goal (MCLG) set by the United States Environmental Protection Agency (EPA).^17,18^ The pesticide has been reported to cause severe health risks when consumed by children, making it an important target molecule for this work.^17–19^

This work builds off of our recent report demonstrating the ability of the LysR-type transcriptional regulator (LTTR) AtzR to detect its cognate ligand, cyanuric acid (CYA), using *in vitro* CFE reactions.^20^ When freeze-dried, the CYA sensor activated only when rehydrated with unfiltered pool water samples containing high (hundreds of micromolar) concentrations of CYA. Here, we combine that CYA cell-free sensor with a reconstituted cell-free metabolic pathway that converts atrazine to CYA through three steps in a single pot reaction. Due to the burden imposed by synthesizing several proteins *in situ* in a single batch CFE reaction, we developed an extract mixing strategy, where individual extracts enriched with a single enzyme or transcription factor are combined to reconstitute the complete biosensing reaction. This modular approach allows the system to be optimized by simply searching over the ratios of each distinct enriched extract. Using this approach, we developed a sensor capable of discriminating high concentrations of atrazine (10-100 μM) within an hour of incubation. We anticipate that our combined metabolism and biosensing strategy for detecting atrazine will be broadly applicable for the rapid cell-free detection of pesticides and other water contaminants, as well as environmental biomarkers and human performance analytes.

## RESULTS

To design our cell-free atrazine sensor, we took inspiration from *Pseudomonas* sp. strain ADP-1, which metabolizes atrazine into cyanuric acid through a three-enzyme pathway encoded on the *atzABC* operon (**Figure 1A**).^21,22^ We hypothesized that by co-expressing each of these enzymes, as well as the cyanuric acid-inducible transcription factor (AtzR) and a fluorescent reporter construct we had previously engineered^20^, we would observe atrazine-inducible reporter protein synthesis. However, co-expressing five different proteins would likely diminish reporter titer and delay the response time for the sensor, even if cell-free expression of each protein were feasible. Instead, we pre-enriched four separate extracts (one each with AtzA, AtzB, AtzC, and AtzR) by inducing protein overexpression from the *E. coli* host strain before lysing the cells and preparing individual extracts. This approach, based on previous work for use in cell-free metabolic engineering,^23^ greatly simplifies the overall sensor design and optimization since the load of each enzyme in the final cell-free reaction can be controlled by adjusting the fraction of its pre-enriched extract in the final mixture. We prepared all of the extracts using methods that maximize gene expression from the endogenous *E. coli* transcriptional machinery.^24^

**Figure 1.**
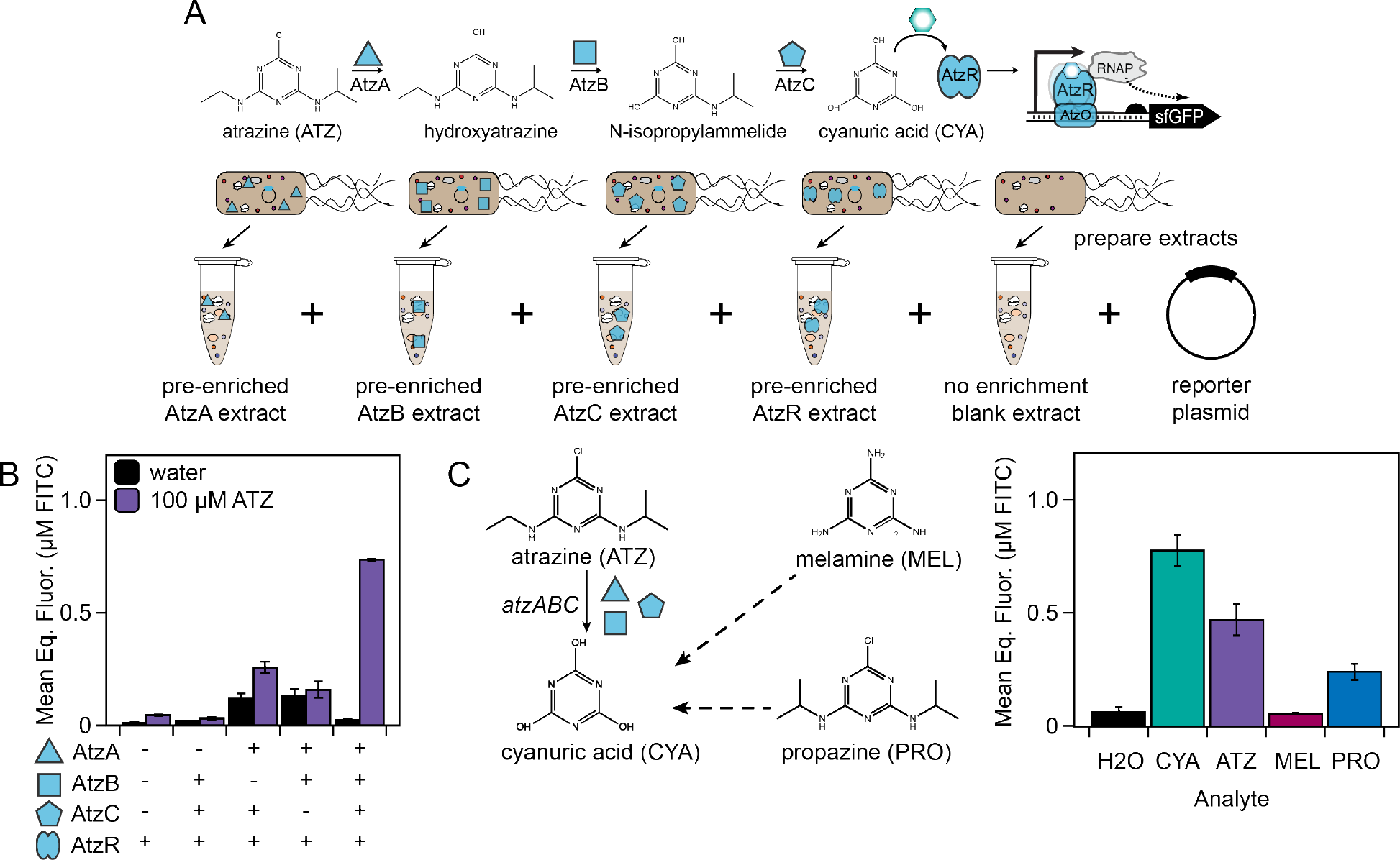
Design of a cell-free atrazine biosensor. (**A**) Atrazine is catabolized by a three-enzyme pathway into cyanuric acid, which can activate transcription through the AtzR transcription factor which recognizes an operator sequence within an engineered promoter. The four required proteins (three enzymes AtzA, AtzB, AtzC and the transcription factor AtzR) are separately overexpressed in strains of BL21 *E. coli*, then lysed and prepared into individual extracts, which can be mixed in fixed ratios to optimize detection of atrazine. (**B**) Cell-free detection of atrazine requires the presence of all three pathway enzymes. (**C**) The sensor is capable of detecting cyanuric acid and atrazine, as well as propazine, a triazine of similar chemical structure, but discriminates against melamine. CYA = cyanuric acid; ATZ = atrazine; MEL = melamine; PRO = propazine. All reactions include 10 nM of a fluorescent reporter DNA template. Error bars represent the standard deviation of sfGFP fluorescence measurements from three technical replicates, correlated to a known linear fluorescein isothiocyanate (FITC) standard.

By mixing all of these protein-enriched extracts together with a “blank” extract that was not pre-enriched with any protein, as well as a reporter plasmid encoding superfolder green fluorescent protein (sfGFP) under regulation by an engineered ADP-1 *atzR* promoter, we could detect atrazine doped into a cell-free reaction at 100 μM (**Figure 1B**), the stoichiometric equivalent of the concentration of cyanuric acid that saturated AtzR activation in our previous work.^20^ As expected, if any of the individual enzyme-enriched extracts was left out of the reaction and replaced with a blank, unenriched extract, the sensor could not effectively detect atrazine above a control reaction supplemented only with water (**Figure 1B**). To the best of our knowledge, this result is the most complex demonstration to date for coupling an upstream metabolic module to an inducible transcriptional biosensor, either in cells or in cell-free systems.

To investigate the sensor’s specificity, we challenged the reaction with two other environmentally relevant triazines that had negligible inhibitory effects on unregulated cell-free transcription and translation reactions (**Supplementary Figure S1**): melamine and propazine. When the biosensor was challenged with each of these compounds, we observed weak activation only in response to propazine, which has a more similar chemical structure to atrazine than melamine (**Figure 1C**). This result is consistent with previous observations that ADP-1’s atrazine chlorohydrolase, AtzA, which actually evolved from the melamine deaminase TriA in the twentieth century, no longer shows any activity on melamine.^25,26^

Satisfied that our cell-free sensor, composed of four different protein-enriched *E. coli* extracts and a blank extract, could detect saturating levels of atrazine, we next aimed to optimize its sensitivity. In the previous mix-and-match approach (**Figure 1B**), we observed a large amount of variability in the negative (water) controls over different extract combinations. This disparity is likely caused both by general batch-to-batch inconsistencies between the extracts^27^, and enzyme-specific poisoning effects, where expression of toxic proteins impacts cell growth and results in poorly performing extracts. To this second point, we found that the protein-enriched extracts were generally less productive than the unenriched extract in a control reaction expressing unregulated sfGFP (**Supplementary Figure S2**). The AtzB and AtzR-enriched extracts, which were prepared from highly growth-inhibited strains, could not support any measurable sfGFP synthesis on their own. However, because of the open reaction environment of the cell-free reaction, we could iteratively adjust enzyme and transcription factor levels by mixing different ratios of the four enriched extracts. To “buffer” the mixed extract against toxicity effects, we made up the rest of the reaction volume with the blank unenriched extract, with the aim of increasing overall reporter protein synthesis yield.

We then aimed to identify the ratio of AtzA-:AtzB-:AtzC-:AtzR-enriched:unenriched extracts that gave the highest fold induction (defined as fluorescence in the presence of 100 μM atrazine / fluorescence in the absence of atrazine) for the sensor. We first determined the optimal fraction of the transcription factor AtzR in the sensing reaction by only detecting the downstream analyte, cyanuric acid. The greatest fold induction was observed at a 5% AtzR-enriched extract and 95% unenriched extract, a ratio that minimizes leak and maximizes ON state, likely because that mixture also has the greatest amount of the unenriched extract (**Figure 2A**). Next, we iteratively optimized the ratios of AtzA-, AtzB-, and AtzC-enriched extracts in the reaction, starting from an assumption of 10% dosage for each sensor (**Figure 2B-D**). Surprisingly, we observed low sensitivity of the atrazine response to perturbations in the concentrations of these enzyme-enriched extracts, at least in the range of 1-10% of the total extract composition. As before, though, no response to atrazine could be observed if any of the enzyme-enriched extracts was individually left out. At each condition, we chose the extract ratio that gave the highest fold induction. Using this iterative, coarse-grained optimization, we obtained an optimal response at 5% AtzR, 10% AtzA, 2% AtzB, and 20% AtzC-enriched extracts, with the balance (63%) made up by the unenriched extract.

**Figure 2.**
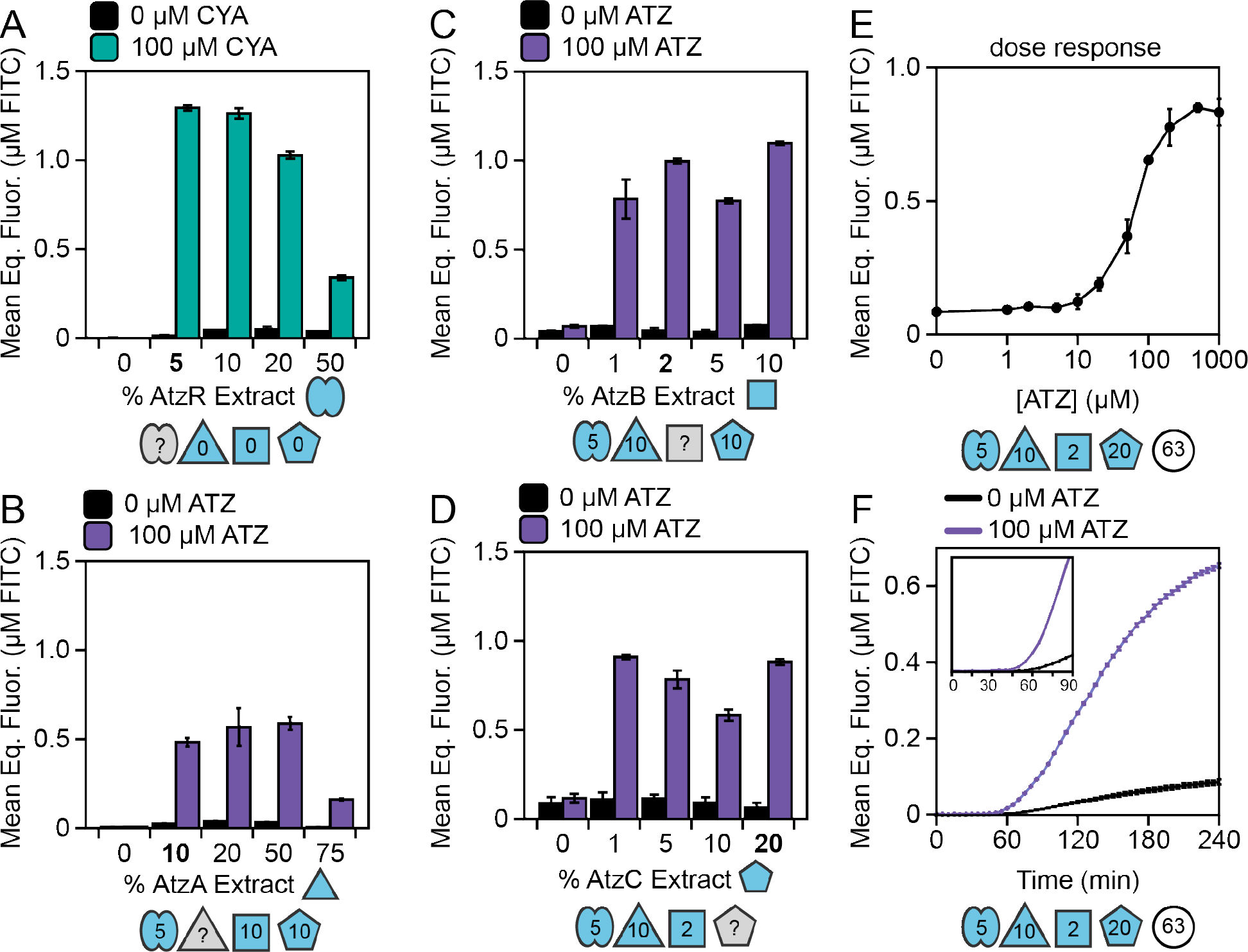
Optimization of the cell-free atrazine sensor. (**A-D**) Iterative optimization of the ratios of the AtzR, AtzA, AtzB, and AtzC extracts in the final sensor mixture reveals that the optimal fold induction is achieved at 5% AtzR-enriched, 10% AtzA-enriched, 2% AtzB-enriched, 20% AtzC-enriched, and 63% unenriched. Symbols below each plot denote the percentage of AtzR (bilobe), AtzA (triangle), AtzB (square), AtzC (pentagon), and blank (circle) extracts included in each experiment. Blank extract was added to make 100% in each condition. Each reaction contained 10 nM of the reporter DNA template from **Figure 1**. (**E**) Atrazine dose-response curve for the optimized sensor, measured by taking defined endpoint samples from biosensor reactions containing the indicated amounts of atrazine, suggests that the overall limit of detection is around 20 μM, which is consistent with previous results reported for the cell-free cyanuric acid sensor.^20^ (**F**) sfGFP fluorescence can distinguish the ON from OFF state within one hour in the presence of 100 μM atrazine, and at endpoint achieves approximately 7-fold induction. Error bars represent the standard deviation of sfGFP fluorescence measurements from three technical replicates, correlated to a known linear FITC standard.

Having established an optimal ratio of each extract in our sensor, we measured its dose response to atrazine by the observed sfGFP fluorescence after 4 hours of reaction (**Figure 2E**). The calculated limit of detection, defined as the concentration of atrazine that yielded a statistically significant detectable signal above background, was approximately 20 μM atrazine (p = 0.01, one-tailed t-test), and the half-maximal signal was observed at around 54 μM by logistic regression. Although these concentrations are far from the stringent limit on atrazine in drinking water set by the EPA, they do approach some of the highest concentrations of atrazine previously detected in raw water (237 ppb = 1.1 μM).^17^ Overall, in response to 100 μM atrazine, the cell-free sensor achieves approximately 7-fold induction and achieves a detectable signal over background in approximately one hour (**Figure 2F**). This work represents an improvement in speed and dynamic range over the previous state-of-the-art whole-cell atrazine biosensor, which reported a similar atrazine sensitivity but suffered from high leak and poor signal-to-noise that would preclude its practical use.^28^

## DISCUSSION

To the best of our knowledge, this work represents the most sensitive genetically encoded biosensor for atrazine. While the sensor is not as sensitive as the 3 ppb atrazine limit set by the EPA, it has significant improvements in speed and dynamic range over previous whole-cell sensors for atrazine.^28^ For multistep enzyme pathways, a cell-free sensor has other practical advantages when compared to a cellular sensor. In particular, we observed severe growth inhibition when overexpressing AtzB (**Supplementary Figure S2**), an effect consistent with previous difficulties in converting a whole-cell cyanuric acid biosensor into one detecting atrazine.^28^ Pre-expressing this protein in a source strain, rather than attempting to make it *in situ*, mitigates these inhibitory effects, especially since the protein expression burden is buffered by a highly productive unenriched extract. Furthermore, our pre-expression approach shifts cell extract resources away from enzyme and transcription factor production and toward reporter synthesis. The resulting higher ON state is another advantage of reconstituting these complex metabolism-sensing pathways *in vitro*. Finally, we believe that our approach of extract mixing, where the dosage of each enzyme or transcription factor is proportional to its fractional composition in the reaction, is an optimal linear tuning strategy, preferable to addressing the nonlinearities introduced by manipulating promoter strength or plasmid copy number, either in cells or in extracts.

Overall, the quantitative agreement of the atrazine transfer function presented in this work to the one we previously demonstrated for cyanuric acid^20^ suggests that this sensor’s limit of detection (LoD) is likely constrained by AtzR’s micromolar affinity for cyanuric acid. A secondary LoD may be set by the similarly low affinity (~50 μM) of the native AtzA for atrazine.^29^ To improve sensitivity to discriminate lower, more environmentally relevant concentrations of atrazine, protein engineering of the transcription factor and enzymes is likely necessary.

More broadly, previous efforts to engineer cell-free systems as molecular biosensors have mainly been limited to well-studied transcriptional regulators like tetracyclines and acyl homoserine lactones.^30,31^ Our strategy of preparing and mixing bacterial extracts pre-enriched with enzymes and transcription factors should greatly expand the scope of molecules that can be detected *in vitro*. The approach can be generalized to sensing any organic molecule that is catabolized, through a pathway of any length, into a metabolite that can be detected by microbes. Since our extract mixing strategy only requires cell-free expression of a single protein, the resource limitations of batch cell-free reactions are minimized.^32^ Thus, cell-free sensors could be multiplexed into a single reaction tube, just by mixing in several more enriched extracts and one new reporter plasmid per sensor.

Overall, this work expands upon previous efforts to build cell-free sensors by detecting an important water contaminant at environmentally relevant levels, and it also proposes a new generalizable approach for building new, highly modular sensors. We anticipate that this design will spur further innovation in cell-free biosensing with an aim toward improving global water security and human health.

## ACKNOWLEDGEMENTS

This work was supported by the Air Force Research Laboratory Center of Excellence for Advanced Bioprogrammable Nanomaterials (C-ABN) Grant FA8650-15-2-5518 (to M.C.J. and J.B.L.), the U.S. Defense Advanced Research Projects Agency’s (DARPA) Living Foundries program award HR0011-15-C-0084, the David and Lucile Packard Foundation (to M.C.J.), the Camille Dreyfus Teacher-Scholar Program (to M.C.J. and J.B.L.), an NSF CAREER Award (Grant 1452441 to J.B.L.) and Searle Funds at the Chicago Community Trust (to J.B.L.). A.D.S. was supported in part by the National Institutes of Health Training Grant (T32GM008449) through Northwestern University’s Biotechnology Training Program.

## MATERIALS AND METHODS

### Plasmid Construction

The reporter plasmid pJBL7030 and AtzR expression plasmid pJBL7032 were used as previously reported.^20^ The AtzA, AtzB, and AtzC-overexpression plasmids pJBL7034, pJBL7035, and pJBL7036 were constructed using Gibson assembly into the pJL1 vector using genes synthesized by Twist Biosciences. All plasmids will be deposited on Addgene with the following IDs: pJBL7034 (133869); pJBL7035 (133870); pJBL7036 (133881); pJBL7030 (133882); pJBL7032 (133883).

### Cell-Free Extract Preparation

Cell-free extract was prepared according to our previously published method.^24^ In general, the host strain for extract preparation was BL21 (DE3) dLacZ. The *lacZ* deficient (lac operon deletion) BL21-Gold-dLac (DE3) strain was a gift from Jeff Hasty (Addgene plasmid # 99247).^9^ The exception to this was the AtzB-enriched extract; since we could not successfully transform the purified plasmid into the knockout strain, this extract was instead prepared using the Rosetta2 (DE3) pLysS strain. For each transformed strain, 1 L culture of 2X YT + P (16 g tryptone, 10 g yeast extract, 5 g NaCl, 7 g potassium phosphate dibasic, 3 g potassium phosphate monobasic) was inoculated from a saturated overnight culture of the chassis strain in LB and grown shaking at 220 RPM at 37°C. For the enzyme- and transcription factor-enriched extract, protein synthesis was induced by adding IPTG at a concentration of 0.5 mM around OD_600_ 0.5. The cells were harvested mainly at OD_600_ 3.0. However, the AtzB and AtzR-enriched extracts were severely growth-restricted, and the cells were harvested once they reached a stationary OD_600_, at approximately 1.5 and OD 2.0 respectively. The remainder of the cell-free preparation followed a published protocol for activating cell-free endogenous transcription,^24^ including an 80-minute runoff reaction and 3-hour dialysis.

### Cell-Free Gene Expression Reaction

Cell-free gene expression reactions were assembled as previously reported from extract, midi-prepped plasmid DNA, salts, nucleotide triphosphates, amino acids, phosphoenolpyruvate, and other cofactors and coenzymes.^24^ 8 mM Mg-glutamate salt solution was used for all experiments. Reactions were assembled in triplicate on ice, and 10 μL of each assembled reaction was pipetted into an individual well of a 384-well plate (Corning, 3712) and measured on a BioTek Synergy H1m plate reader. sfGFP fluorescence was measured using emission at 485 nm and excitation at 520 nm. Fluorescence values were correlated to equivalents of fluorescein isothiocyanate (FITC) using a known standard curve.

### Data Availability

Source data for all figures and the FITC standard curve are available in the Northwestern University Arch Institutional Repository (doi 10.21985/n2-z5vp-tk94).

