## Supplementary Information for "Design and optimization of a cell-free atrazine biosensor"

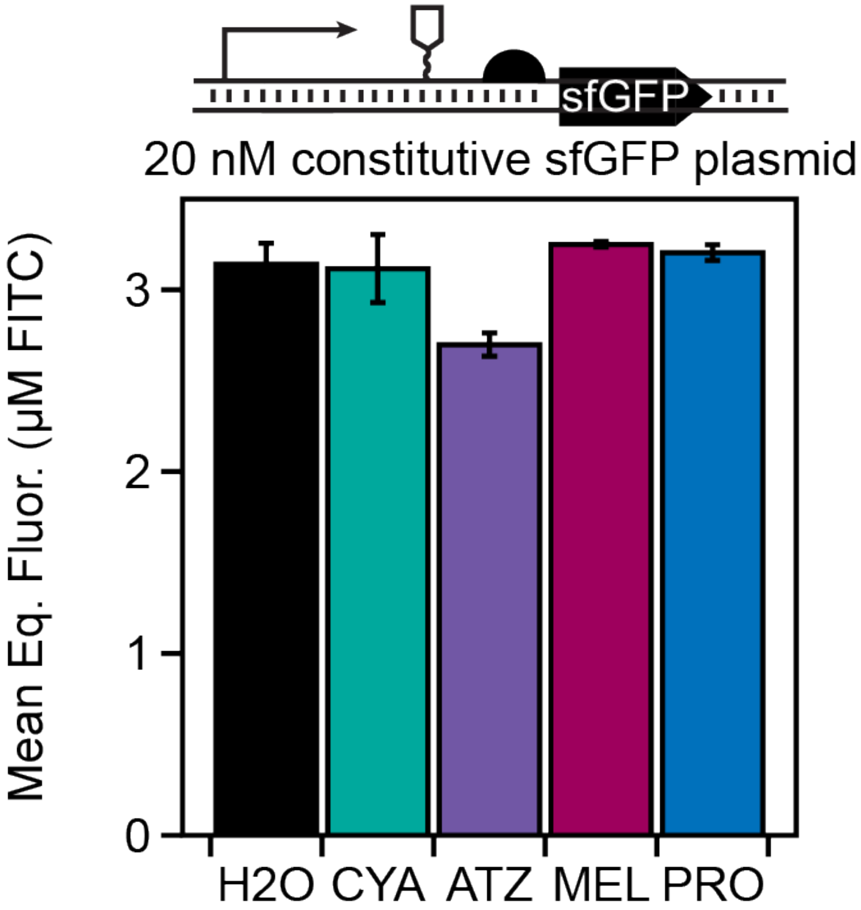


**Supplementary Figure 1. Investigating potential triazine poisoning of unregulated CFE reactions.** A constitutively expressed sfGFP reporter plasmid was included in a cell-free expression reaction supplemented with 100 µM of the triazines atrazine, melamine, or propazine to investigate potential poisoning effects. The plasmid used for this experiment was the previously reported pJBL7030, which has the J23119 promoter (consensus $\sigma^{70}$ -10 and -35 recognition sites) and a pHP14 stability hairpin. Only atrazine minimally inhibits the gene expression reaction. Error bars represent the standard deviation of sfGFP fluorescence measurements, correlated to a known linear FITC standard, from three technical replicate reactions. CYA = cyanuric acid; ATZ = atrazine; MEL = melamine; PRO = propazine.

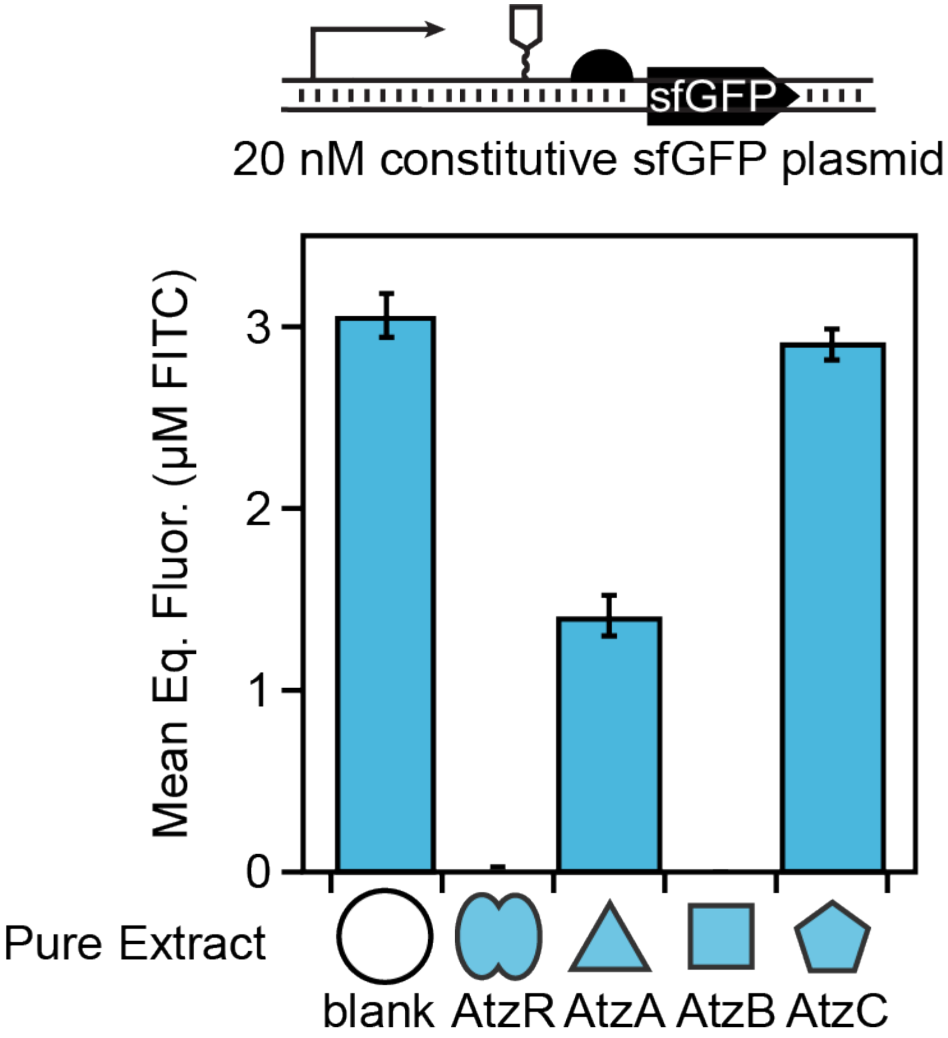


**Supplementary Figure 2. Poising of extracts enriched for specific biosensor components**. Unenriched extract (blank) and extracts enriched for AtzR, AtzA, AtzB and AtzC were each used to express an unregulated sfGFP reporter plasmid (pJBL7030). Significant poisoning effects can be observed from extracts enriched with AtzR, AtzA and AtzB. Overexpression of AtzR and AtzB also resulted in severe growth defects in the host strain, consistent with the resulting low extract yield. These results motivated an extract mixing strategy where hybrid extract performance can be rescued by the addition of a blank unenriched extract. Error bars represent the standard deviation of sfGFP fluorescence measurements, correlated to a known linear FITC standard, from three technical replicate reactions.
